# Altered active control of step width in response to mediolateral leg perturbations while walking

**DOI:** 10.1101/2019.12.30.890947

**Authors:** Nicholas K. Reimold, Holly A. Knapp, Rachel E. Henderson, Landi Wilson, Alyssa N. Chesnutt, Jesse C. Dean

**Author notes:** Corresponding author information: Jesse Dean, 77 President St., Charleston, SC 29425-5712 MSC700, USA.

## Abstract

During human walking, step width is predicted by mediolateral motion of the pelvis, a relationship that can be attributed to a combination of passive body dynamics and active sensorimotor control. The purpose of the present study was to investigate whether humans modulate the active control of step width in response to a novel mechanical environment. Participants were repeatedly exposed to a force-field that either assisted or perturbed the normal relationship between pelvis motion and step width, separated by washout periods to detect the presence of potential after-effects. As intended, force-field assistance directly strengthened the relationship between pelvis displacement and step width. This relationship remained strengthened with repeated exposure to assistance, and returned to baseline afterward, providing minimal evidence for assistance-driven changes in active control. In contrast, force-field perturbations directly weakened the relationship between pelvis motion and step width. Repeated exposure to perturbations diminished this negative direct effect, and produced larger positive after-effects once the perturbations ceased. Both of these results provide evidence of gradual changes in active control in response to perturbations. In the longer term, these methods may be useful for improving deficits in the active control of step width often observed among clinical populations with poor walking balance.

## Introduction

In human walking, mediolateral motion of the pelvis during a step predicts step width at the end of the step. This behavior has long been cited as important for ensuring mediolateral walking balance [1], as a “dynamically-appropriate step width” that accounts for pelvis motion would presumably be wide enough to prevent a lateral loss of balance toward the stepping leg, but not so wide as to cause excessive mediolateral velocities in the opposite direction during the next step. Indeed, larger mediolateral pelvis displacements and velocities away from the stance foot have been linked to wider steps, as recently reviewed by Bruijn and van Dieen [2]. This relationship can be mathematically quantified using partial derivatives, regressions, or correlations [3-4], consistently revealing a significant relationship between step width and pelvis motion throughout a step.

However, the existence of statistically significant correlations between pelvis motion and step width is alone not sufficient to prove that this relationship is being actively controlled by the nervous system. The passive dynamics of the body (e.g. segment inertial properties) likely play an important role, as simply allowing the stance leg and torso to act as an inverted pendulum could contribute to the observed relationship. Recent simulations have demonstrated that at least in the sagittal plane, passive dynamics can produce significant correlations between pelvis motion and foot placement without need for within-step active control [5]. This limitation of correlation-based methods has been widely acknowledged [2-3, 6], and has motivated the use of other experimental methods to provide more direct evidence for active control of step width. Namely, pelvis dynamics early in a step predicts the magnitude of within-step swing phase hip abductor activity, which in turn predicts mediolateral foot placement location [6]. A similar pattern of active control is seen with the application of mechanical perturbations, as perturbations that increase the pelvis mediolateral displacement or velocity away from the stance leg elicit increased swing phase hip abductor activity and more lateral foot placement [6-8]. Sensory perturbations that create the perception of an altered mechanical state provide further evidence for active control, as perturbations of either hip proprioception [9-10], vision [11] or vestibular feedback [12] are followed by changes in foot placement location consistent with the previously described mechanical principles [3].

While substantial evidence indicates a role for active control in the step-by-step adjustments of step width, it is presently unclear whether humans readily modulate this control to meet the demands of the environment. As a step toward investigating potential changes in active control, we have developed a novel elastic force-field able to push users’ legs toward targeted step widths [13]. Walking in this force-field thus involves an additional contributor to the step-by-step relationship between pelvis motion and step width, as schematically illustrated in Figure 1. In our initial work, we found that controlling the force-field to produce minimal mediolateral forces on users (Transparent mode) [13] had a negligible effect on the relationship between pelvis motion and step width [14]. We also used the force-field to “assist” participants toward a dynamically-appropriate step width, based on the step-by-step motion of their pelvis. This assistive approach had the direct effect of increasing the strength of the relationship between pelvis motion and step width, as quantified using the partial correlation between step width and pelvis displacement at the start of the step (step start ρ_disp_) [14]. Conversely, we also used the force-field to “perturb” participants, decreasing the strength of the relationship between pelvis motion and step width [14]. Ceasing assistance was followed by short-lived after-effects in which the link between pelvis motion and step width was weakened, while ceasing perturbations caused after-effects in which this link was strengthened [14].

**Figure 1.**
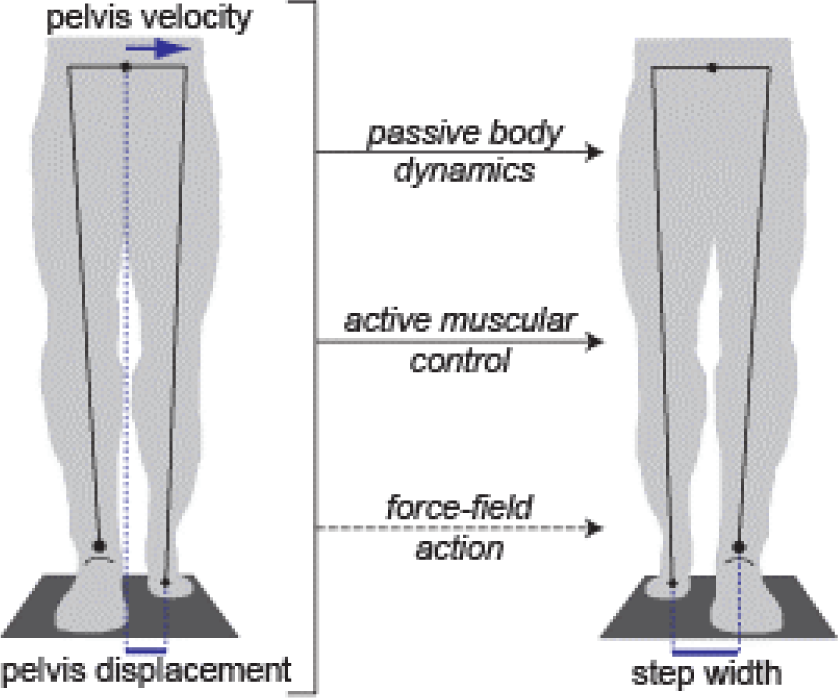
Mediolateral pelvis displacement and velocity at the start of a step (left panel) predict step width at the end of the step (right panel), here illustrated in a frontal plane view of a step with the left foot. This relationship can be influenced by frontal plane passive body dynamics, as the swing leg could act as a pendulum, and the stance leg as an inverted pendulum. Active muscular control can also contribute to this relationship, for instance by using the swing leg hip abductors to influence swing leg position, and the stance leg hip abductors to influence motion of the pelvis relative to the stance leg. In the present study, our force-field is an additional contributor to this relationship, as it can be used to influence step width based on pelvis displacement at the start of the step.

Extending our prior results, the purpose of the present study was to investigate changes in the within-step active control of step width upon repeated exposure to an altered mechanical environment produced by our force-field. While the relationship between pelvis motion and step width can be influenced by multiple factors (Fig. 1), our primary comparisons focused on periods in which the force-field action (and passive dynamics of the body) remained constant. Therefore, changes in the relationship between pelvis motion and step width can be attributed to altered active control. Similar approaches of varying mechanical context and quantifying changes in gait kinematics over time have been used extensively to investigate altered control in other novel environments, including split-belt walking [15-17], walking with leg swing assistance or resistance [18], and walking with leg joint trajectories assisted or perturbed [19-20]. Across these distinct contexts, initial exposure to the novel environment often causes an immediate large change in the movement pattern (here termed a *direct effect*), which gradually decays back toward baseline as active control is adjusted with repeated experience moving in this environment. Subsequent return to the original mechanical environment often also causes an immediate change in the movement pattern (here termed an *after-effect*), followed by a return to baseline as active control is again adjusted. With repeated exposure to the same novel environment, the magnitude of both the direct effects and after-effects can decrease, likely as individuals more rapidly adjust their active control to the change in environment [21-23].

We investigated changes in the step-by-step relationship between pelvis motion and step width that can be attributed to altered active control, testing four specific hypotheses. First, we hypothesized that upon initial exposure to an altered mechanical environment, the immediate direct effects of the force-field will decrease over time, as participants adjust their active control. Second, we hypothesized that upon the initial removal of the altered mechanical environment, the magnitude of after-effects will decrease toward the baseline level over time. Third, we hypothesized that repeated exposure to the altered mechanical environment will reduce the direct effects produced by the force-field. Finally, we hypothesized that with repeated exposure, the after-effects observed with removal of the force field will decrease. Each of these hypotheses was tested for both force-field assistance and perturbations. As the major focus of this study was changes in the within-step control of step width, our primary measure was step start ρ_disp_, just as in our prior work [14]. Changes in this correlation would indicate that active control of this within-step relationship has been altered. Secondarily, we quantified the partial correlation between step width and pelvis displacement at the end of the step (step end ρ_disp_). This metric quantifies the extent to which any within-step adjustments influence the relationship between pelvis state and step width once the new base of support is established, as is likely important for mediolateral balance [2]. Finally, our secondary analyses also included the more traditional gait metrics of step width and step length.

## Results

### Gait changes during an initial Transparent trial

While the purpose of this study was to investigate gait changes in response to an altered mechanical environment, we first performed a control comparison of whether gait behavior changed over the course of a 5-minute Transparent trial (with minimal mediolateral forces) *before* participants were exposed to either force-field assistance or perturbations. The relationship between mediolateral pelvis motion and step width did not vary from early to late in the first Transparent trial. Specifically, neither step start ρ_disp_ (p=0.72; 0.67±0.13 early vs. 0.66±0.13 late; mean±s.d.) nor step end ρ_disp_ (p=0.98; 0.79±0.09 early vs. 0.79±0.10 late) differed between the early and late periods. In contrast, both step width (p<0.001) and step length (p<0.001) varied significantly over the course of the trial, with step width decreasing (150±29 mm early vs. 136±30 mm late) and step length increasing (645±33 mm early vs. 654±30 mm late).

### Gait changes spanning three force-field exposures

For illustrative purposes, we here depict the changes in each gait metric across three exposures to force-field assistance or perturbations, interspersed with washout periods in Transparent mode. The clearest effects on step start ρ_disp_ (Fig. 2a) and step end ρ_disp_ (Fig. 2b) were observed during the periods in which assistance or perturbations were applied, with assistance increasing and perturbations decreasing ρ_disp_. During the washout periods, the ρ_disp_ metrics returned toward their baseline level (dashed line), sometimes exhibiting overshoot beyond this value. In general, step width tended to decrease over time (Fig. 2c), while step length increased (Fig. 2d). The increases in step width and decreases in step length relative to baseline were more apparent during the periods with assistance than those with perturbations.

**Figure 2.**
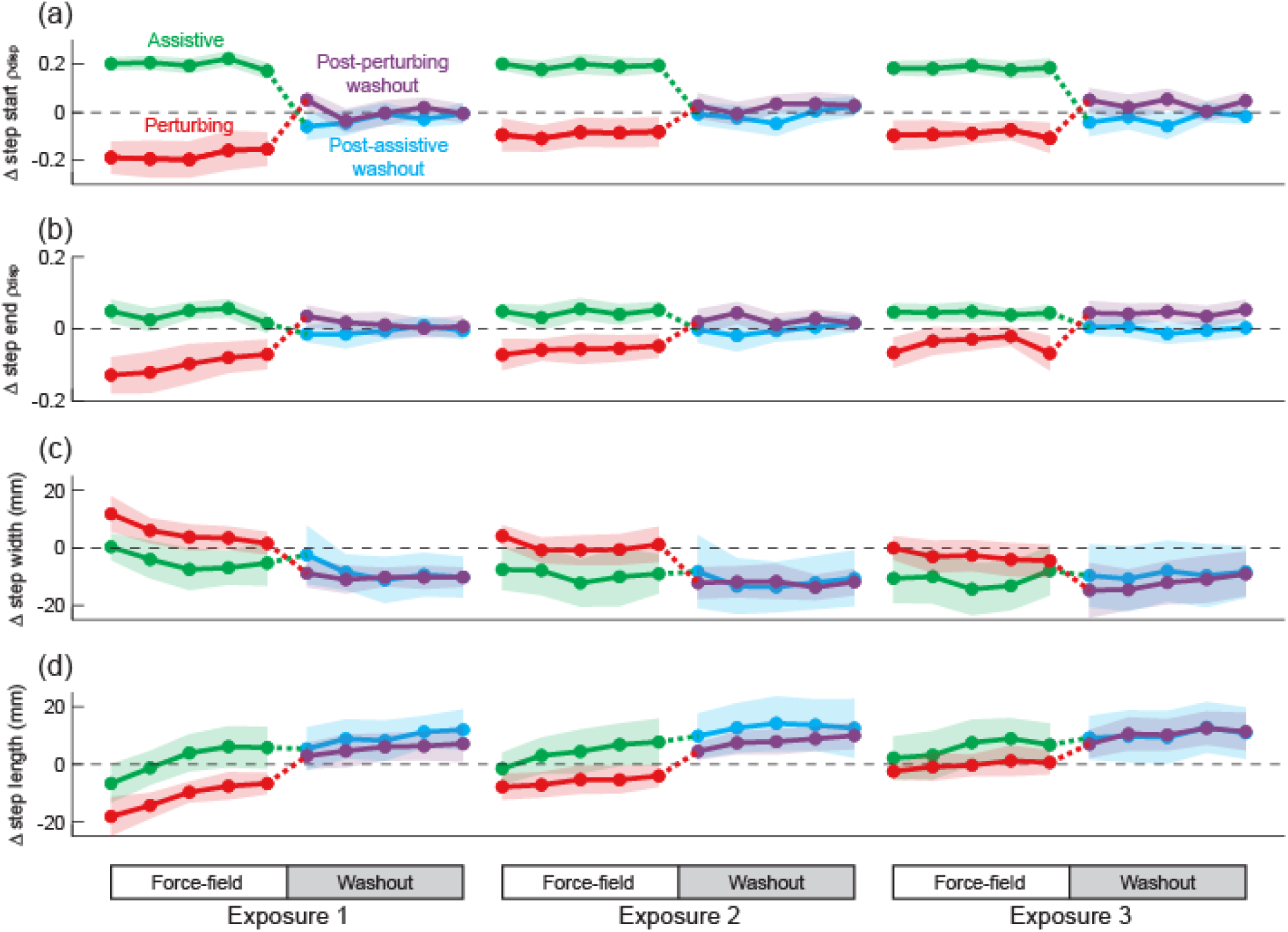
Gait behavior across three consecutive 10-minute walking trials (exposures) that included either force-field assistance or perturbations. These changes are illustrated for step start ρ_disp_ (a), step end ρ_disp_ (b), step width (c), and step length (d), and are plotted in terms of the difference from the initial Transparent trial. Each data point represents the mean value calculated within each 45-step bin, and shaded areas indicate 95% confidence intervals. The dashed line at zero is presented to allow easier visualization of changes relative to the initial Transparent trial.

### Direct effects during the first force-field exposure

Force-field assistance increased the strength of the relationship between pelvis displacement and step width (ρ_disp_) throughout a step (Fig. 3a). Over the duration of the first exposure to the assistance, step start ρ_disp_ decreased significantly (p=0.015; Fig. 3b), while remaining higher than the baseline Transparent level. Conversely, we did not observe a significant change in step end ρ_disp_ (p=0.063; Fig. 3c). Step width did not change (p=0.13; Fig. 3d) across the initial exposure to force-field assistance, while step length increased significantly (p=0.002; Fig. 3e).

**Figure 3.**
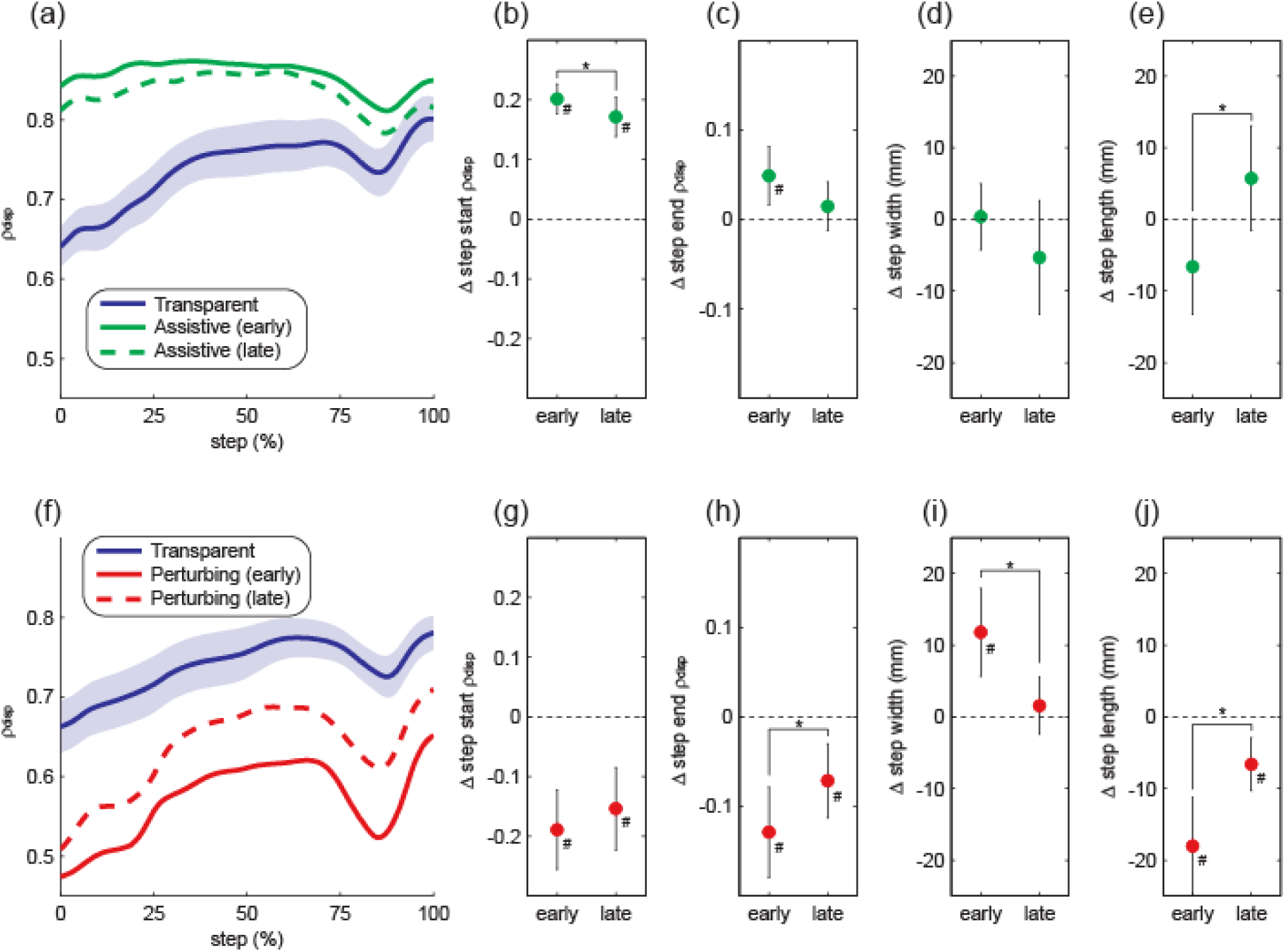
Direct effects of the first force-field exposure. The top row illustrates the direct effects of force-field assistance, in terms of ρ_disp_ magnitude throughout the step (a) and our four gait outcome measures (b-e). The bottom row (f-j) follows the same structure to illustrate the direct effects of force-field perturbations. In panels (a) and (f), the shaded area indicates the 95% confidence interval for the initial Transparent trial, while confidence intervals are not shown for the experimental conditions to avoid extensive overlap. For the remaining panels, data are presented as the difference from the initial Transparent trial. Data points indicate means and error bars indicate 95% confidence intervals. Asterisks (*) indicate a significant difference between the indicated early and late periods. Pound signs (#) indicate a significant difference from the initial Transparent trial, with the 95% confidence interval not including zero (dashed line).

Opposite to the effects observed with force-field assistance, perturbations weakened the relationship between pelvis displacement and step width throughout a step (Fig. 3f). This decrease in step start ρ_disp_ did not change over the course of this initial trial (p=0.36; Fig. 3g), while step end ρ_disp_ increased significantly closer to its baseline value (p=0.018; Fig. 3h). Both step width (p<0.001; Fig. 3i) and step length (p=0.001; Fig. 3j) changed significantly over the course of the perturbation trial, with step width decreasing and step length increasing.

### After-effects during the first washout period

Following the first exposure to force-field assistance, the magnitude of ρ_disp_ throughout the step did not vary from early to late in the subsequent washout period (Fig. 4a). No significant differences were observed for step start ρ_disp_ (p=0.14; Fig. 4b) or step end ρ_disp_ (p=0.55; Fig. 4c) between these time periods. However, step width decreased significantly (p=0.016; Fig. 4d) and step length increased significantly (p=0.012; Fig. 4e) over this same period.

**Figure 4.**
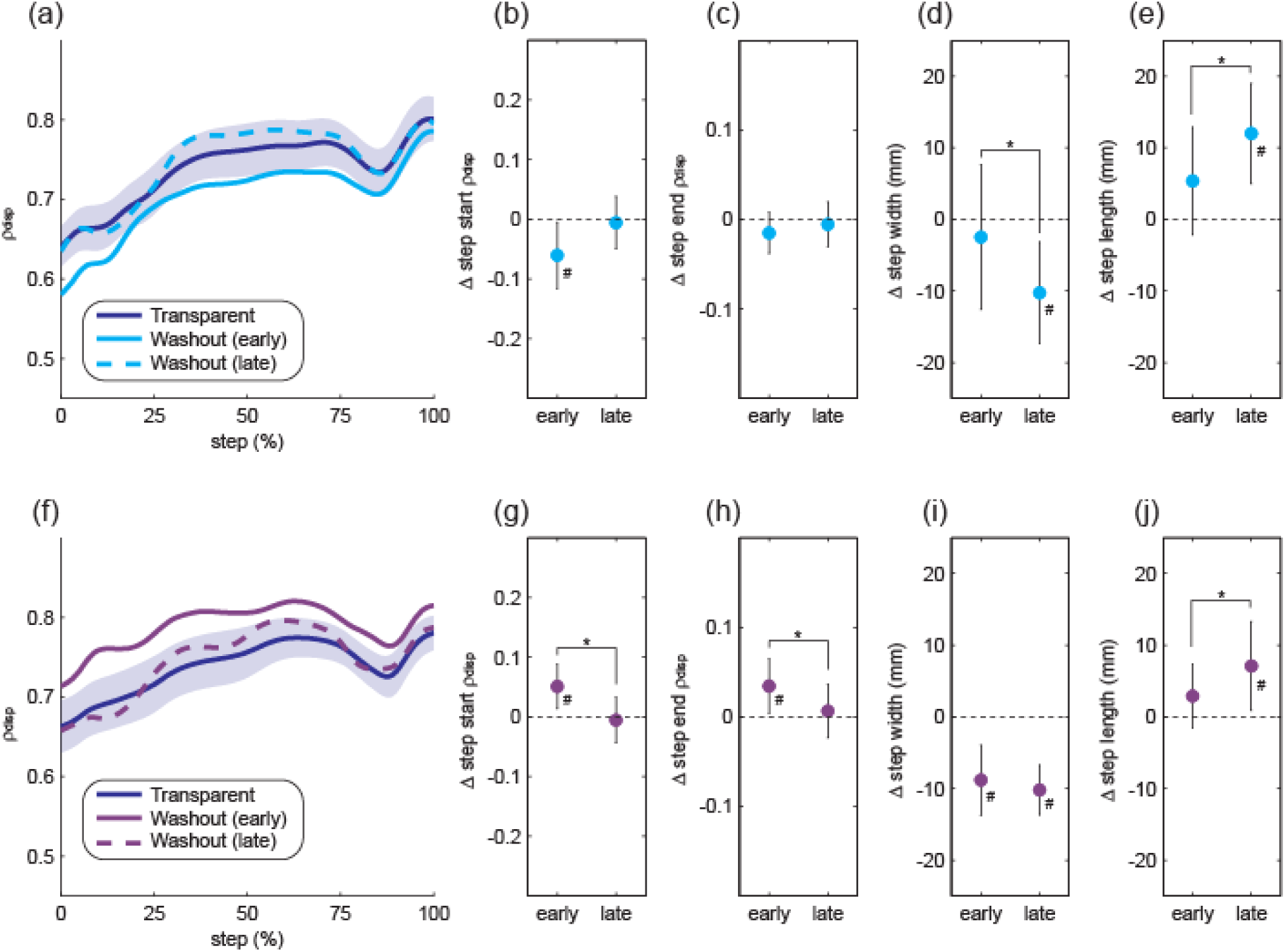
Gait behavior over the course of the first washout period. The top row (a-e) illustrates potential after-effects of force field assistance, while the bottom row (f-j) illustrates potential after-effects of force-field perturbations. The structure of the figure is the same as that described for Figure 3.

Over the course of the first washout period after force-field perturbations, ρ_disp_ magnitude decreased significantly toward its baseline value (Fig. 4f), as observed for step start ρ_disp_ (p=0.030; Fig. 4g) and step end ρ_disp_ (p=0.040; Fig. 4h). For both of these metrics, ρ_disp_ magnitude did not differ from the baseline Transparent trial by late in the washout period. Step width did not change (p=0.52; Fig. 4i) from early to late in the first washout trial, while step length increased significantly (p=0.009; Fig. 4j).

### Direct effects with repeated force-field exposure

Across all three exposures to force-field assistance, ρ_disp_ magnitude throughout the step remained elevated relative to its baseline value (Fig. 5a). No significant differences were observed across these exposures in terms of direct effects on step start ρ_disp_ (p=0.27; Fig. 5b) or step end ρ_disp_ (p=0.96; Fig. 5c). Step width decreased significantly across these exposures (p=0.009; Fig. 5d), while step length did not change (p=0.56; Fig. 5e).

**Figure 5.**
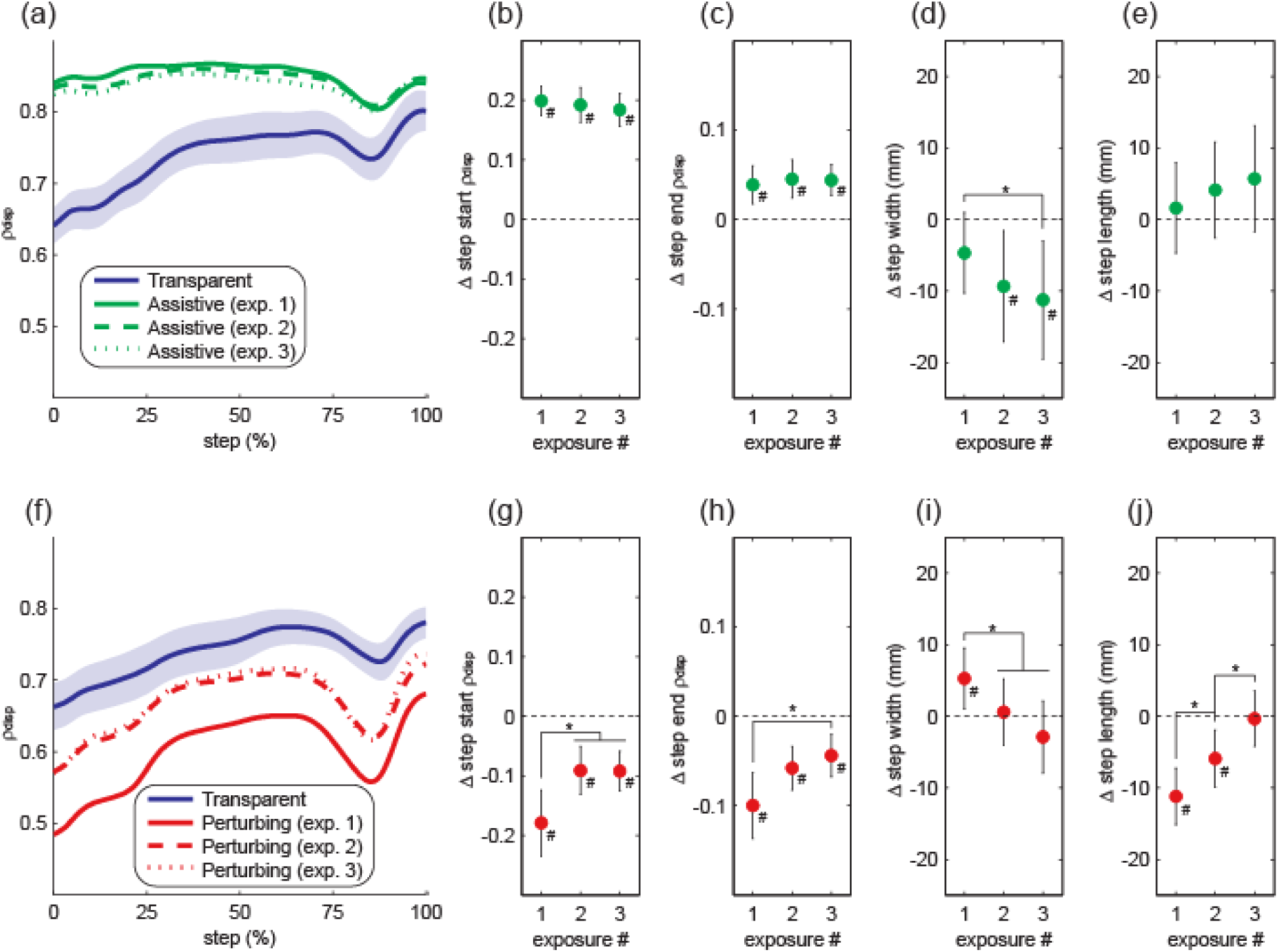
Effects of repeated exposure to the force-field. Following the structure of Figure 3, the top row (a-e) illustrates the direct effects of force-field assistance, while the bottom row (f-j) illustrates the direct effects of force-field perturbations. Asterisks (*) indicate significant differences between the indicated exposures. Pound signs (#) indicate a significant difference from the initial Transparent trial.

Changes in the direct effects of repeated force-field exposure were more apparent with perturbations. The ρ_disp_ magnitude throughout the step was decreased relative to its baseline value for all exposures but was closer to baseline during the second and third exposures (Fig. 5f). For both step start ρ_disp_ (p=0.044; Fig. 5g) and step end ρ_disp_ (p=0.030; Fig. 5h), the direct effects of perturbations were significantly smaller in later exposures. Repeated exposure also significantly influenced the direct effects of perturbations on step width (p<0.001; Fig. 5i) and step length (p<0.001; Fig. 5j), as step width decreased and step length increased in later exposures.

### After-effects with repeated force-field exposure

The observed ρ_disp_ magnitudes throughout the step were similar during each of the washout periods following force-field assistance (Fig. 6a). No significant differences were present between these washout periods in terms of step start ρ_disp_ (p=0.58; Fig. 6b) or step end ρ_disp_ (p=0.75; Fig. 6c). Similarly, no significant differences were present between the washout periods for step width (p=0.26; Fig. 6d) or step length (p=0.56; Fig. 6e).

**Figure 6.**
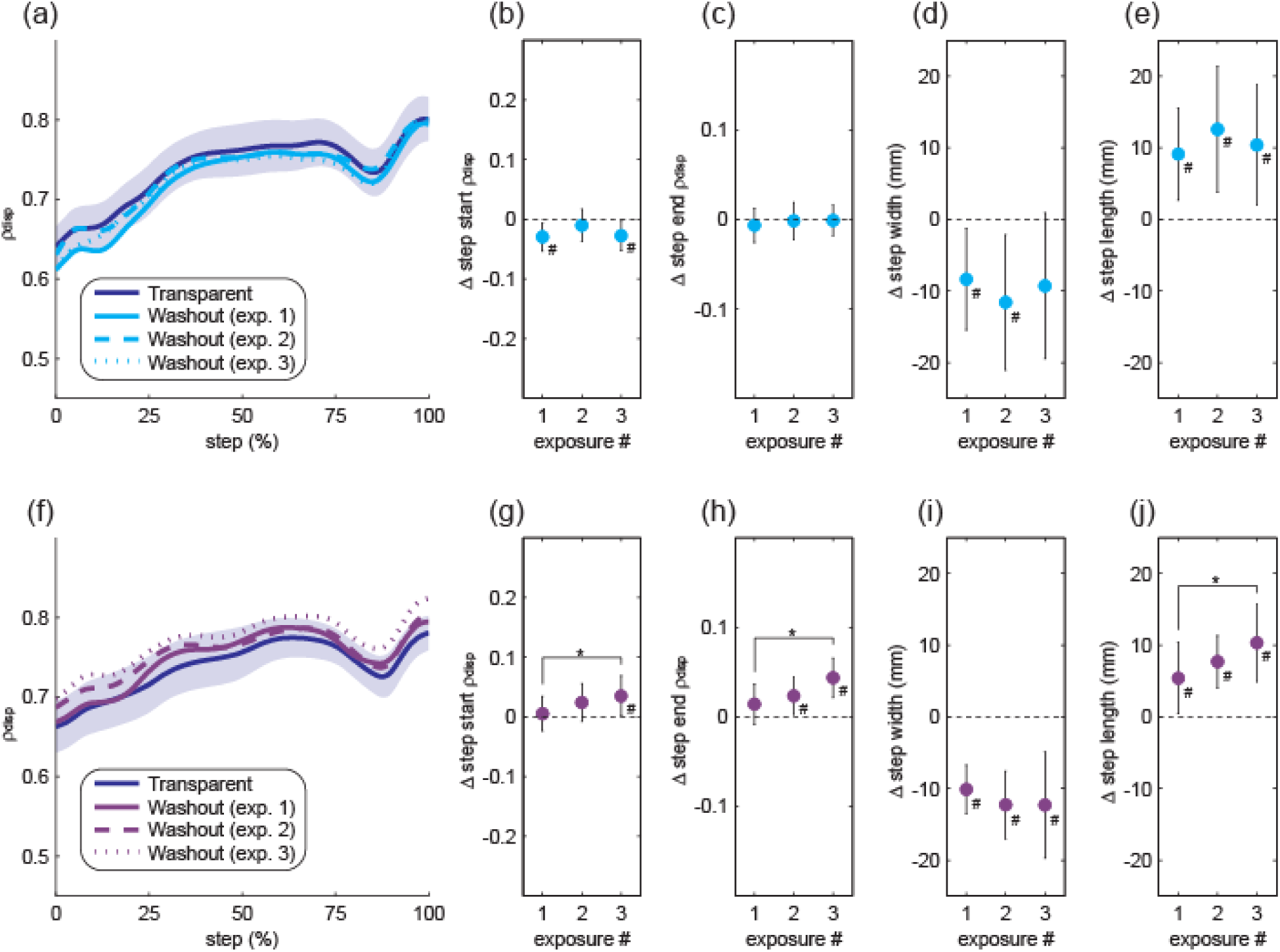
Gait behavior across the three washout periods. The top row (a-e) again illustrates gait metrics during the three washout periods following force-field assistance, while the bottom row (f-j) illustrates these metrics during washout periods following perturbations. The structure of the figure follows Figure 5.

During the washout periods following force-field perturbations, ρ_disp_ magnitudes throughout the step increased for later exposures (Fig. 6f). Both step start ρ_disp_ (p=0.042; Fig. 6g) and step end ρ_disp_ increased significantly (p=0.014; Fig. 6h) following later perturbation exposures. Step width did not vary across consecutive washout periods (p=0.17; Fig. 6i), but step length increased significantly (p=0.017; Fig. 6j) in later washout periods.

## Discussion

Exposure to a novel force-field caused changes in the step-by-step relationship between mediolateral pelvis motion and step width. While both force-field assistance and perturbations had measurable effects on this relationship, our hypotheses regarding altered active control within a consistent mechanical context were only partially supported. In general, we observed stronger evidence for altered active control in response to perturbations than in response to assistance. Both the positive direct effects of assistance and the positive after-effects of perturbations were retained with repeated exposure to the force-field, a finding of interest for the potential future application of these methods as a clinical intervention.

While the present study was not a direct replication of our previous force-field experiments, the changes in gait behavior observed upon initial exposure to assistive or perturbing forces were consistent with our prior results. Specifically, we previously found that force-field assistance (across varied control equations) directly strengthened the link between pelvis motion and step width, as quantified with ρ_disp_ at the start of the step [14]. Upon cessation of the assistance, we observed short-lived after-effects in which step start ρ_disp_ was decreased relative to baseline. In contrast, force-field perturbations weakened the link between pelvis motion and step width, and caused subsequent short-lived positive after-effects [14]. The same patterns of direct effects and after-effects on step start ρ_disp_ were seen in the present study upon the first exposure to assistance or perturbations.

Here, our focus was on changes in gait behavior across extended periods of walking while the force-field remained in the same control mode (i.e. Transparent, Assistive, Perturbing). Our primary metric quantified the relationship between pelvis displacement at the start of the step and step width. While multiple factors can contribute to the strength of this within-step relationship (including passive dynamics) [2-3, 5-6], comparing periods in which the mechanical context remained the same allows us to attribute changes to altered active control. Secondarily, we quantified the relationship between pelvis displacement at the end of the step and step width, providing insight into the ultimate effect of any within-step adjustments. This metric is conceptually similar to step-by-step variability in the mediolateral margin of stability calculated from the body’s extrapolated center of mass [3], a common measure of walking balance. Importantly, neither of these metrics (step start ρ_disp_ and step end ρ_disp_) changed from early to late in the initial Transparent trial, indicating that any observed changes are not simply due to altered active control with extended periods of walking.

Exposure to force-field assistance produced only weak evidence for changes in active control, with qualitatively similar effects for step start ρ_disp_ and step end ρ_disp_. Over the course of the first exposure to assistance, ρ_disp_ tended to decrease toward its baseline value, although this decrease did not reach significance for step end ρ_disp_. The reduction in step start ρ_disp_ may be a result of the assistance allowing participants to safely exert less active control over this within-step relationship – a “slacking” phenomenon previously observed with assistive robotic orthoses [24]. However, the hypothesized changes in ρ_disp_ during the first washout period and with repeated exposure to assistance were not directly supported by our results. Across all three exposures, ρ_disp_ remained elevated while assistance was applied, and essentially returned to baseline when the assistance ended. The lack of large changes in ρ_disp_ across repeated bouts may be due to the assistive nature of the forces, as mechanical contexts that *challenge* walking balance appear more likely to cause adjustments in active control than contexts not perceived as challenging [25]. The present results also provide no evidence for changes in active control that would cause the altered movement pattern (with increased ρ_disp_) to be retained once the assistance ends, consistent with prior results in which force-fields were used to assist achievement of a specific footpath trajectory during walking [19-20].

Force-field perturbations were often accompanied by changes in ρ_disp_ indicative of altered active control. The direct effect of perturbations was to weaken the link between pelvis displacement and step width; however, this relationship tended to return toward its baseline level with extended or repeated periods of exposure. Similar results have been observed with force-fields that perturb the footpath trajectory during walking [20, 26]. While we did not observe a significant increase in step start ρ_disp_ from early to late in the first perturbation exposure, this may have simply been due to the relatively short (5-minute) duration of this first exposure. Indeed, changes in gait kinematics during spit-belt walking can continue to develop across five 15-minute exposures on consecutive days [23], suggesting that the adjustment of active control may require an extensive period of time [27]. Speculatively, we attribute the gradual reduction in the effects of force-field perturbations over repeated exposures to adjustments in the active control used to resist these perturbations, which may otherwise increase the risk of a lateral loss of balance. Changes in ρ_disp_ during the initial washout period following perturbations provide further evidence for altered active control, with such after-effects previously observed with other perturbing force-fields [20, 26], and commonly cited as a strong indicator of changes in sensorimotor control [17]. The relationship between pelvis displacement and step width was strengthened early in the first washout period but returned to its baseline level – consistent with a return to the initial pattern of active control. However, the changes in ρ_disp_ during washout periods following repeated perturbation exposures contradicted our hypothesis. Instead of observing decreased after-effects with repeated exposure, step start ρ_disp_ and step end ρ_disp_ actually increased significantly. This unexpected result is perhaps due to the fact that the altered active control during these washout periods produces a stronger link between pelvis motion and step width, and thus is unlikely to have a negative effect on balance that would drive further adjustments in active control.

Although step width and step length often varied both within and across walking trials, we are unable to attribute these changes to the novel mechanical environments produced by the force-field. In general, step width decreased and step length increased over time, changes observed even in the first Transparent trial before assistive or perturbing forces were applied. We suspect that these general trends are not due to the force-field itself, but instead reflect a gradual shift away from a ‘cautious’ gait pattern with short, wide steps [28-31] as participants became accustomed to walking while interfaced with the force-field. As most easily seen in Figure 2, the clearest difference between the effects of the Assistive and Perturbing modes was the wider and shorter steps used while perturbations were delivered. This behavior is likely an example of a ‘generalized anticipatory strategy’ used by participants to maintain their balance [32] in response to a context perceived as potentially destabilizing.

While the present results provide evidence for altered active control of step width when perturbed, a limitation of this work is our inability to identify the underlying physiological mechanism. One possibility is error-based sensorimotor adaptation, in which movement patterns gradually change in response to new mechanical demands [33]. This adaptation is thought to be driven by sensory prediction errors [34], as humans seek to reduce the difference between the predicted and sensed movement through a trial-and-error process. In the present work, force-field perturbations could conceivably increase the errors between the intended and actual step width, causing participants to adjust the active muscle contractions typically used to influence swing leg motion [6]. This altered active control could reduce the observed effects of perturbations, as well as produce the tighter link between pelvis motion and step width once the perturbations cease. Unfortunately, calculating our primary outcome measure (ρ_disp_) requires numerous consecutive steps [4], which prevents us from mimicking analytical approaches from split-belt walking that quantify error-based adaptation using a small number of steps (or even single steps). For example, the largest magnitude after-effects are observed for approximately the first five strides following a change in mechanical environment [35], with the magnitude of such effects influenced by the time course [22], amplitude [36], and pattern [37] of the perturbations. Additionally, the time course of adaptation is often calculated based on changes in step characteristics on a step-by-step basis [21]. Beyond these limitations of our analyses, our observation of increased after-effects with repeated perturbation exposure would not be predicted from error-based adaptation, in which repeated exposure to a novel environment is expected to result in smaller direct and after-effects [21-23]. Perhaps this altered gait behavior would be better explained by the framework of reinforcement learning, in which humans learn new movement patterns based on their perceived value [17]. The tighter link between pelvis displacement and step width may be perceived as valuable for the combined goals of avoiding a lateral loss of balance and not wasting mechanical energy with excessively wide steps. These potential explanations are speculative, and carefully designed future work will be required to differentiate between the potentially contributing mechanisms. Additionally, our finding that perturbations influence the active control of step width may be specific to perturbations that target the legs, as recent work has reported that mediolateral trunk perturbations have a more notable effect on control of the trunk [38].

A further limitation of the present work was the sample size. Participant recruitment was based on our previous study [14], in which this sample size (n=12 per group) was sufficient to detect significant direct effects and after-effects of force-field assistance and perturbations. However, the present study’s statistical comparisons of changes in ρ_disp_ within a given mechanical context resulted in numerous p-values between 0.01 and 0.10, which should be interpreted with caution. Just as the present results supported the primary findings of our original work, future work should attempt to replicate the present results with a larger sample size, based on effect size estimates we can now calculate.

A long-term goal of this line of research is to apply our force-field methods to clinical populations with deficits in walking balance. We have previously observed that the step-by-step relationship between pelvis displacement and step width is significantly weaker for paretic steps than for non-paretic steps among chronic stroke survivors [39], who often have an increased fall risk [40]. Additionally, the links between pelvis displacement, hip abductor activity, and mediolateral foot placement are weakened in stroke survivors with clinically-identified balance deficits [41], providing evidence for altered active within-step control. Paralleling the present results in neurologically-intact controls, future work will test whether similar effects of force-field assistance and perturbations are observed in patients with a reduced link between pelvis motion and step width. The potential to strengthen the link between pelvis motion and step width may serve as a useful intervention for improving post-stroke walking balance. Supporting the feasibility of this approach, repeated exposure to an altered mechanical environment has proven successful in normalizing step length in neurologically-injured populations over extended periods of time, whether using a split-belt walking paradigm [42-43] or mechanical resistance of leg swing [44].

In conclusion, this study provides initial evidence for altered within-step active control of step width in response to targeted force-field perturbations. With extended periods of exposure to these perturbations, participants exhibited an increased ability to both resist the direct effects of the perturbations and maintain the strengthened link between pelvis motion and step width once the perturbations ceased. The apparent effects of force-field assistance on active control were more modest; participants generally continued to “accept” the provided assistance with repeated exposure, but did not exhibit positive after-effects once the assistance ceased. Future work is needed to investigate the potential motor adaptation or learning mechanisms underlying these results, and to test whether similar methods can produce beneficial effects in clinical populations with deficits in walking balance.

## Methods

### Participants

This experiment involved 24 young, neurologically-intact participants who had not previously interacted with the force-field. Participants were randomly assigned to either the Assistive group (n=12; age = 22±1 yrs; height = 170±9 cm; mass = 70±13 kg; mean±s.d.) or the Perturbing group (n=12; age = 23±1 yrs; height = 170±8 cm; mass = 65±13 kg; mean±s.d.). All participants provided written informed consent using a document approved by the Medical University of South Carolina Institutional Review Board, and consistent with the Declaration of Helsinki.

### Force-field Design and Control

We used a custom-designed force-field to exert mediolateral forces on participants’ legs while walking. This force-field has previously been described in detail [13] and used to both assist and perturb the relationship between mediolateral pelvis motion and step width [14]. Pairs of linear actuators (UltraMotion; Cutchoge, NY, USA) positioned anterior and posterior to a treadmill were used to rapidly adjust the mediolateral location of two steel wires running parallel to the treadmill belts, and in series with extension springs. These wires passed through leg cuffs worn on the lateral shank, which allowed free anteroposterior and vertical leg motion. Participants experienced mediolateral leg forces that were proportional to the mediolateral displacement between the actuator end point and the leg cuff [13]. The force-field’s mediolateral stiffness (ratio between mediolateral force and displacement) was 180 N/m, based on the results of a pilot study in which this stiffness was sufficient to produce clear effects on step width (Appendix A).

Three force-field control modes were applied: Transparent, Assistive, and Perturbing. For all modes, actuator positions were controlled based on the location of active LED markers (PhaseSpace; San Leandro, CA, USA) on the sacrum, heels, and/or leg cuffs. The sacrum marker was used to estimate mediolateral pelvis location, a simplification previously found to have minimal effects on our calculations [4]. In Transparent mode, each actuator followed the mediolateral motion of the corresponding leg cuff, minimizing the resultant mediolateral leg forces. In both the Assistive and Perturbing modes, we first used the mediolateral displacement of the pelvis from the stance heel at the start of each step to predict a dynamically-appropriate step width. For example, if a right step began with the pelvis located relatively far to the right of the left stance heel, we would predict that the upcoming step should be relatively wide. This prediction was based on previously collected empirical data quantifying the step-by-step relationship between pelvis displacement and step width [4]. In the Assistive mode, we positioned the actuators to push the swing leg toward the predicted dynamically-appropriate step width. This was done using the following equation, in which *SW* represents the predicted step width, *x*_*pelvis*_ represents the mediolateral displacement of the pelvis from the stance heel at the start of the step, and *SW*_*mean*_ represents the participant’s mean step width:

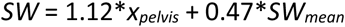

Conversely, in the Perturbing mode, we positioned the actuators to push the swing leg away from the predicted step width (e.g. to encourage a narrow step when a wide step would be dynamically appropriate). This was accomplished using the following equation:

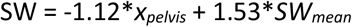

The effects of these force-field control equations were first investigated in our prior work [14], along with several other equations of varying complexity. While all previously tested control equations had similar effects, we observed the largest positive after-effects following force-field perturbations with the equation applied here [14].

### Experimental Procedure

Participants performed five treadmill walking trials at 1.2 m/s, in non-randomized order. All participants wore a harness attached to an overhead rail that did not support body weight but would have prevented a fall in case of a loss of balance. Participants first walked for 5-minutes without interacting with the force-field (Normal), followed by a 5-minute trial with the force-field in Transparent mode. Data from the final 2-minutes of the Normal trial were used to calculate each participant’s mean step width, with this value integrated into the relevant Assistive or Perturbing control equation. Participants in the Assistive group then performed three identical 10-minute trials in which the force-field was in Assistive mode for the first 5-minutes and Transparent mode for the final 5-minutes as a washout period. Participants in the Perturbing group performed three corresponding 10-minute trials in which the force-field was in Perturbing mode for the first 5-minutes and Transparent mode for the final 5-minutes, again as a washout period.

### Data Collection and Processing

Active LED marker locations were sampled at 120 Hz and low-pass filtered at 10 Hz. We defined each step start as the time point when the ipsilateral heel velocity changed from posterior to anterior [45]. The step end was defined as the time point when the contralateral heel velocity changed from posterior to anterior. Throughout each step, we quantified the mediolateral displacement of the pelvis relative to the stance heel, as well as the mediolateral pelvis velocity. Step width was defined as the mediolateral displacement between the ipsilateral heel marker at the step end and the contralateral heel marker at the step start. Step length was calculated as the difference between the anterior position of the ipsilateral heel at the step end and the anterior position of the contralateral heel at the previous step end, accounting for treadmill speed.

Each 5-minute walking period within a given force-field mode was divided into five “bins” for the steps taken with each leg: steps 1-45; steps 46-90; steps 91-135; steps 136-180; steps 181-225. While the number of steps taken within each 5-minute period varied fairly widely across participants and conditions (275±12; mean±s.d.), every participant took at least 225 steps with each leg for each 5-minute period. Comparisons across bins thus included the same number of steps (interactions with the novel mechanical environment), while the inclusion of 45 consecutive steps allowed convergence of our correlation-based measures (detailed below) [4]. An alternative approach of dividing walking trials into bins based on time (i.e. minutes 1-5) produced similar results (Appendix B).

Our primary outcome measure was the partial correlation between mediolateral pelvis displacement and step width (ρ_disp_), accounting for variation in mediolateral pelvis velocity [14]. The strength of the relationship between pelvis motion and step width can alternatively be quantified using the R^2^ magnitude calculated by performing a linear regression between these gait variables [4]. Such R^2^-based analyses essentially parallel the changes in ρ_disp_ (Appendix C), likely because predictions of step width seem to be dominated by pelvis displacement during steady-state walking [4].

For illustrative purposes, we calculated ρ_disp_ over the course of a step for each 45-step bin. We resampled pelvis displacement and velocity values during each step to create 101-sample vectors, thus allowing comparisons across steps of variable periods. We created a trajectory of ρ_disp_ values by calculating ρ_disp_ from the pelvis displacement and velocity values at each normalized time point in the step (from 0-100) and the step width values at the end of the step. While we calculated ρ_disp_ throughout the step, our statistical analyses focused on ρ_disp_ at the start and end of the step, as well as step width and step length.

### Statistics

Upon visual inspection, the gait metrics of interest appeared consistent with a normal distribution. However, we chose to apply non-parametric statistical methods due to the relatively small sample size in this study [46]. We first performed a series of Wilcoxon signed-rank tests (α=0.05) comparing several gait metrics (step start ρ_disp_, step end ρ_disp_, step width, step length) between early (steps 1-45) and late (steps 181-225) in the initial 5-minute Transparent trial, including all 24 participants. Our subsequent statistical tests were focused on specific hypotheses, and were performed separately for participants in the Assistive and Perturbing groups. To test whether the direct effects of the initial force-field exposure changed over time, we performed Wilcoxon signed-rank tests (α=0.05) to compare our gait metrics (step start ρ_disp_, step end ρ_disp_, step width, step length) between early (steps 1-45) and late (steps 181-225) in the first exposure to the Assistive/Perturbing force-field. To test whether potential after-effects that follow the initial force-field exposure changed over time, we performed identically structured Wilcoxon signed-rank tests (α=0.05) to compare these gait metrics between early (steps 1-45) and late (steps 181-225) in the Transparent washout period that followed the first exposure to Assistance/Perturbations. To test whether repeated exposure to the altered mechanical environment influenced the direct effects of the force-field, we performed Friedman’s tests (α=0.05) to compare these gait metrics (combined across all 5 bins) between the three exposures to the Assistive/Perturbing force-field. Finally, to test whether repeated exposure to the altered mechanical environment influenced the magnitude of any after-effects, we performed identical Friedman’s tests (α=0.05) to compare these gait metrics between the three washout periods in Transparent mode. In the case of a significant effect for the Friedman’s test, we performed Wilcoxon signed-rank tests (α=0.017 to account for multiple comparisons) to detect significant differences between individual conditions.

## Author Contributions

JD conceived of the research. NR created software used in data collections. AC, RH, HK, NR, and LW conducted data collections. NR, LW, and JD contributed to data analysis. JD drafted the initial manuscript. All authors reviewed and revised the manuscript.

## Additional Information / Competing Interests Statement

The authors declare no competing interests.

## Acknowledgements

This work was supported by a grant from the National Science Foundation (1603391).

## Data Availability Statement

The datasets generated during the current study are available from the corresponding author on reasonable request.

