## Supplementary Material for "Altered active control of step width in response to mediolateral leg perturbations while walking"

### Appendix A

We performed a pilot experiment in which we varied the effective mediolateral stiffness of the force-field, defined as the ratio between the mediolateral force acting on the leg and the mediolateral displacement between the leg cuff and the actuator end point. The mediolateral stiffness was varied by adjusting the initial tension of the extension spring in series with the wires running parallel to the treadmill belts, as described previously [1].

Ten young, neurologically intact individuals (9 F / 1 M; age = 24±5 yrs; height = 167±10 cm; mass = 64±12 kg; mean±s.d.) participated in this study. None of these participants had previously interacted with the force-field, and all participants provided written informed consent using a document approved by the Medical University of South Carolina Institutional Review Board, and consistent with the Declaration of Helsinki.

All walking trials were performed at 1.2 m/s, and participants wore a harness attached to an overhead rail that did not support body weight, but would have prevented a fall in case of a loss of balance. Participants first performed a 5-minute Normal walking trial in which they did not interface with the force-field. They then performed a series of six 7-minute walking trials, in which the force-field was in either Assistive or Perturbing mode (identical to in the main text) for 5-minutes, followed by a 2-minute washout period with the force-field in Transparent mode. Each participant performed both an Assistive trial and a Perturbing trial (in randomized order) at each of three mediolateral stiffness values (also in randomized order): 60 N/m; 120 N/m; and 180 N/m.

As in the main text, our primary outcome measure was the partial correlation between mediolateral pelvis displacement and step width ($\rho_{disp}$). We calculated $\rho_{disp}$ magnitude throughout the step, while our statistical analyses focused on this metric at the start of the step and the end of the step. Additionally, for this pilot experiment our analyses focused only on the direct effects of the force-field, during the periods in which the force-field was in either Assistive or Perturbing mode. Mimicking our main text analyses, we compared step start $\rho_{disp}$ and step end $\rho_{disp}$ combined across the first five consecutive 45-step bins during these 5-minute walking periods.

To test whether mediolateral stiffness influenced the direct effects of force-field assistance, we performed a Friedman's test ($\alpha$=0.05) to compare step start $\rho_{disp}$ and step end $\rho_{disp}$ across the three stiffness values. The same statistical analysis structure was used to test whether mediolateral stiffness influenced the direct effects of force-field perturbations. In the case of a significant effect for the Friedman's test, we performed Wilcoxon signed-rank tests ($\alpha$=0.017 to account for multiple comparisons) to detect significant differences between individual conditions.

*Effects of mediolateral force-field stiffness.* As may be expected, stiffer force-fields generally produced larger effects on the gait pattern. With force-field assistance, higher stiffnesses appeared to produce larger increases in $\rho_{disp}$ throughout the step relative to baseline (Fig. A1a). This difference was statistically significant for step start $\rho_{disp}$ ($p<0.001$; Fig. A1b), but not for step end $\rho_{disp}$ ($p=0.24$; Fig. A1c). With force-field perturbations, higher stiffnesses produced larger decreases in $\rho_{disp}$ throughout the step (Fig. A1d). This larger negative effect for higher stiffnesses was observed for both step start $\rho_{disp}$ ($p=0.011$; Fig. A1e) and step end $\rho_{disp}$ ($p<0.001$; Fig. A1f).

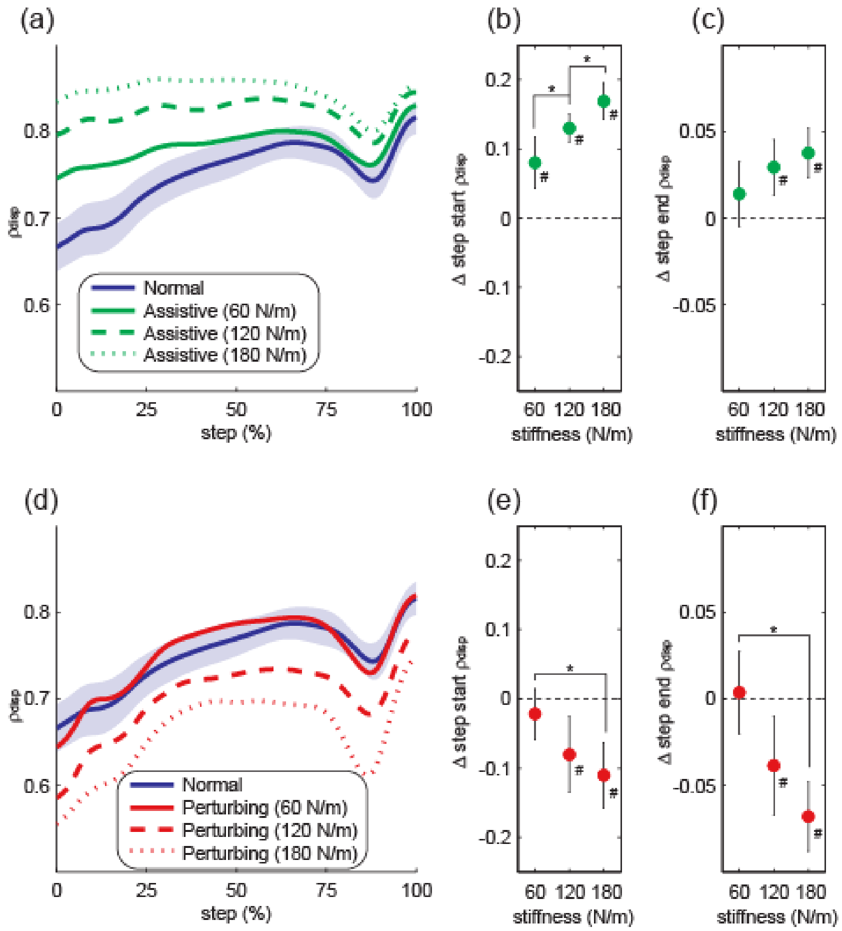


**Figure A1.** Influence of force-field stiffness on the direct effects of force-field assistance and perturbations. The top row illustrates the direct effects of force-field assistance on $\rho_{disp}$ magnitude throughout the step (a), step start $\rho_{disp}$ (b), and step end $\rho_{disp}$ (c). The bottom row (d-f) follows the same structure to illustrate the direct effects of force-field perturbations. In panels (a) and (d), the shaded area indicates the 95% confidence interval for the initial Normal trial. For the remaining panels, data are presented as the difference from the initial Normal trial. Data points indicate means and error bars indicate 95% confidence intervals. Asterisks (*) indicate a significant difference between the indicated stiffness values. Pound signs (#) indicate a significant difference from the initial Normal trial, with the 95% confidence interval not including zero (dashed line).

### Appendix B

In the main text, we divided each 5-minute walking period into bins consisting of 45 consecutive steps. As each step can be considered a single interaction with the novel mechanical environment created by the force-field, this approach allowed us to perform comparisons across individual participants based on the same number of interactions. Here, we illustrate the same comparisons as in Figure 2 of the main text, but instead divide the walking periods into 1-minute bouts.

The same primary effects seen in the main text are also present with this time-based method, as illustrated in Figure B1. Briefly, the clearest effects on step start $\rho_{disp}$ (Fig. B1a) and step end $\rho_{disp}$ (Fig. B1b) are again observed during the periods in which assistance or perturbations were applied, as assistance increased and perturbations decreased these metrics. During washout periods, these metrics returned to their baseline level, sometimes exhibiting overshoot. Again, step width tended to decrease over time (Fig. B1c), and step length tended to increase over time (Fig. B1d).

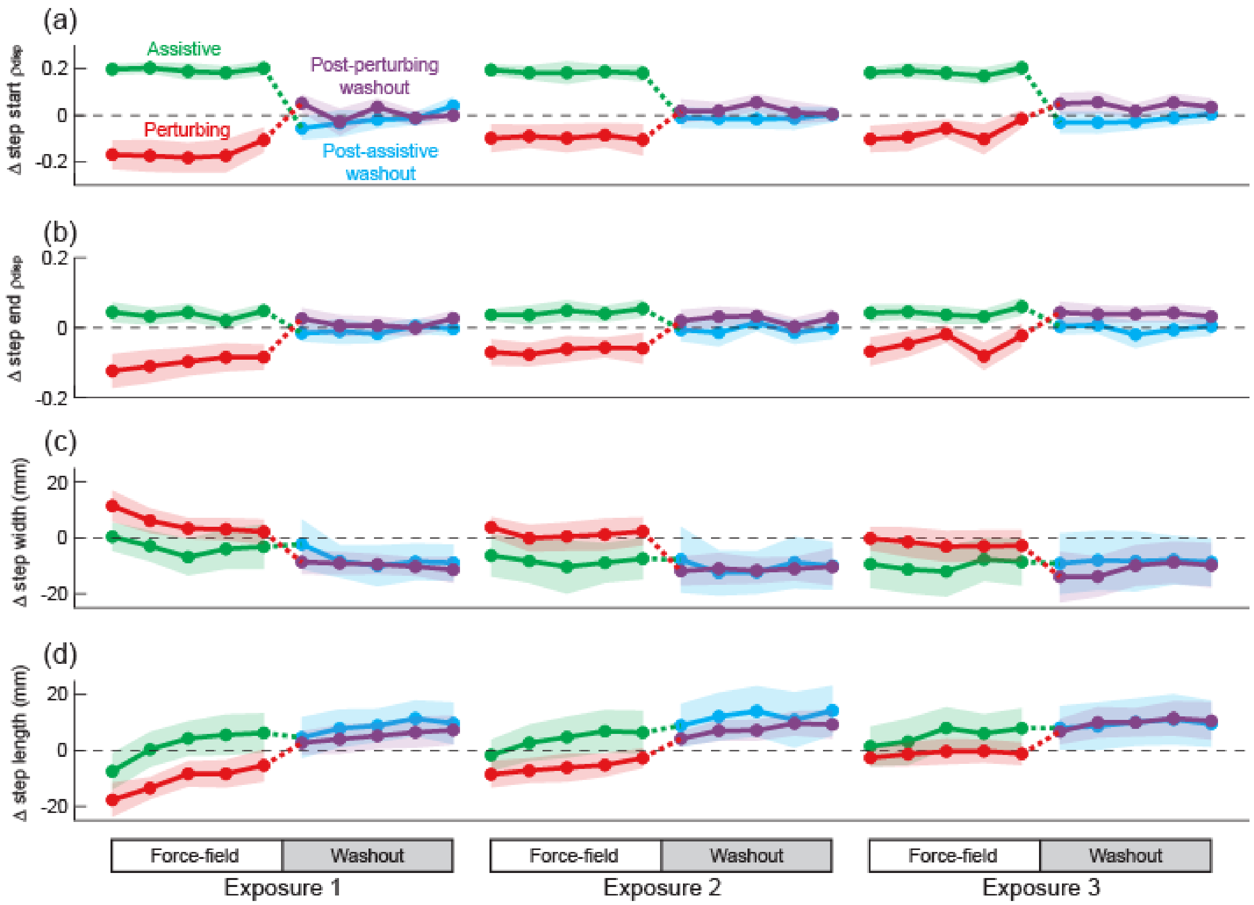


**Figure B1.** Gait behavior varied over three consecutive 10-minute walking trials that included either force-field assistance or perturbations. These changes are illustrated for step start $\rho_{disp}$ (a), step end $\rho_{disp}$ (b), step width (c), and step length (d), and are plotted in terms of the difference from the initial Transparent trial. Each data point represents the mean value calculated within each minute of walking, and shaded areas indicate 95% confidence intervals. The dashed line at zero is presented to allow easier visualization of changes relative to the initial Transparent trial.

### Appendix C

As in previous work [2-3], we here performed linear regressions with mediolateral pelvis displacement and velocity as independent variables, and step width as the dependent variable. The $R^2$ value resulting from this regression was used to quantify the strength of the relationship between pelvis motion and step width. As with our primary analyses focused on $\rho_{disp}$, we calculated $R^2$ magnitude based on pelvis motion throughout the step. Our statistical analyses focused on step start $R^2$ and step end $R^2$, mirroring our analyses of $\rho_{disp}$ in our main text. Overall, the results seen with $R^2$ were quite similar to those presented for $\rho_{disp}$ in the main text, as detailed below.

*Direct effects during first force-field exposure.* Force-field assistance increased $R^2$ magnitude throughout the step, both early and late in the first assistance exposure (Fig. C1a). Over the course of this initial exposure, step start $R^2$ decreased significantly ($p=0.028$; Fig. C1b), while step end $R^2$ did not ($p=0.61$; Fig. C1c). In contrast, force-field perturbations decreased $R^2$ magnitude throughout the step, with these decrements observed both early and late in the first perturbation exposure (Fig. C1d). Over the course of this initial exposure, step start $R^2$ did not change ($p=0.30$; Fig. C1e), while step end $R^2$ increased significantly toward its baseline level ($p=0.014$; Fig. C1f).

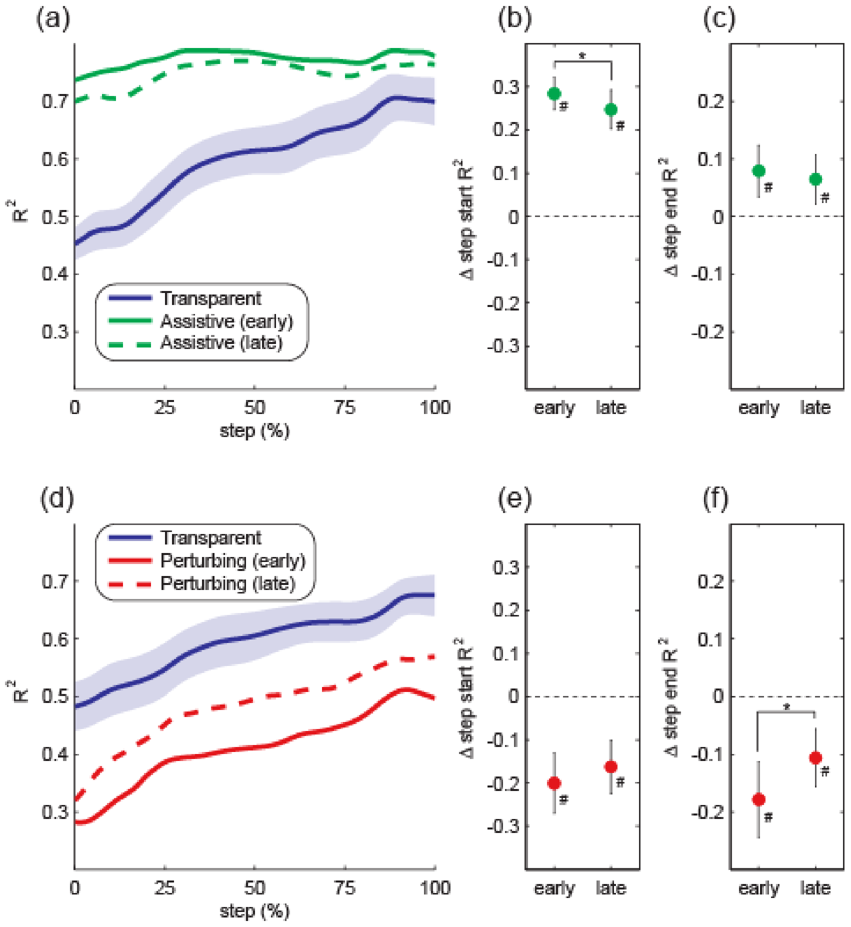


**Figure C1.** Direct effects of the first force-field exposure. The top row illustrates the direct effects of force-field assistance on $R^2$ magnitude throughout the step (a), step start $R^2$ (b), and step end $R^2$ (c). The bottom row (d-f) follows the same structure to illustrate the direct effects of force-field perturbations. In panels (a) and (d), the shaded area indicates the 95% confidence interval for the initial Transparent trial. For the remaining panels, data are presented as the difference from the initial Transparent trial. Data points indicate means and error bars indicate 95% confidence intervals. Asterisks (*) indicate a significant difference between the indicated early and late periods. Pound signs (#) indicate a significant difference from the initial Transparent trial, with the 95% confidence interval not including zero (dashed line).

*After-effects during first washout period*. Following the first exposure to force-field assistance, $R^2$ magnitude throughout the step did not vary significantly from early to late in the subsequent washout period (Fig. C2a). No significant differences were observed for step start $R^2$ ($p=0.19$; Fig. C2b) or step end $R^2$ ($p=0.55$; Fig. C2c) between these time periods. Over the course of the first washout period after force-field perturbations, $R^2$ magnitude appeared to decrease toward its baseline value (Fig. C2d). This decrease from early to late in the washout period was significant for step start $R^2$ ($p=0.043$; Fig. C2e), but not for step end $R^2$ ($p=0.063$; Fig. C2f).

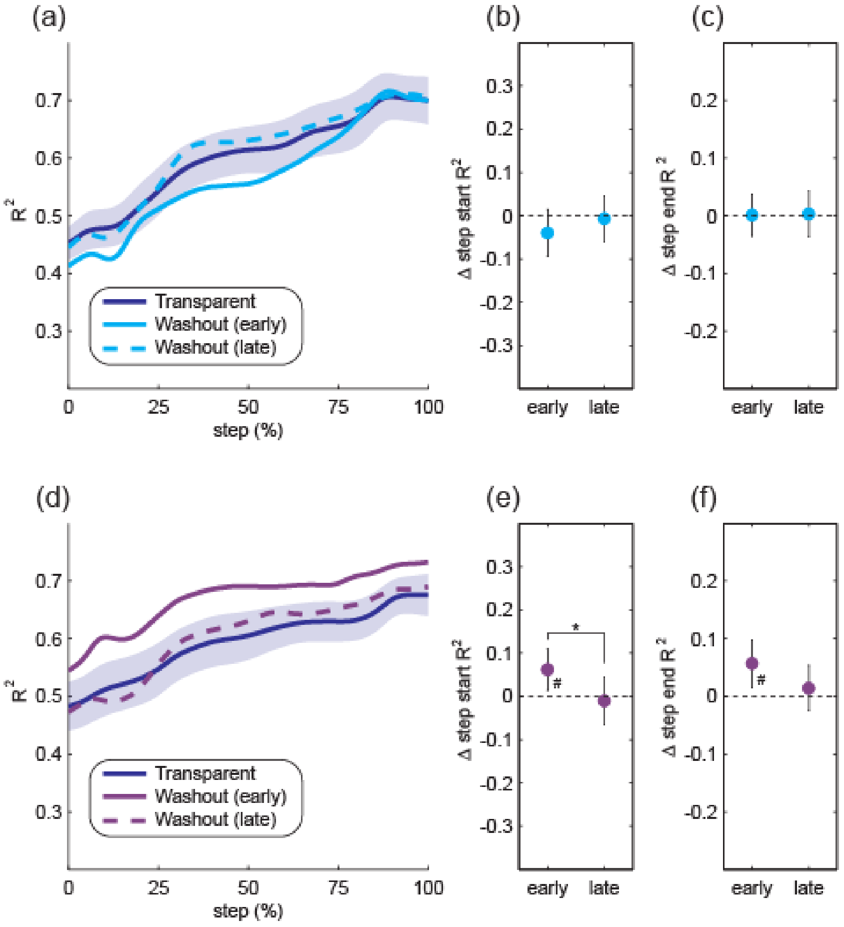


**Figure C2.** Gait behavior over the course of the first washout period. The top row (a-c) illustrates potential after-effects of force field assistance, while the bottom row (d-f) illustrates potential after-effects of force-field perturbations. The structure of the figure is the same as that described for Figure C1.

*Direct effects with repeated force-field exposure.* Across the three exposures to force-field assistance, $R^2$ magnitude was increased throughout the step (Fig. C3a). Neither step start $R^2$ ($p=0.09$; Fig. 3b) nor step end $R^2$ ($p=0.75$; Fig. C3c) changed significantly across exposures, remaining elevated relative to baseline. While $R^2$ magnitude throughout the step was decreased with force-field perturbations, this metric returned toward its baseline value with repeated exposure (Fig. C3d). Both step start $R^2$ ($p=0.011$; Fig. C3d) and step end $R^2$ ($p<0.001$; Fig. C3e) changed significantly across exposures.

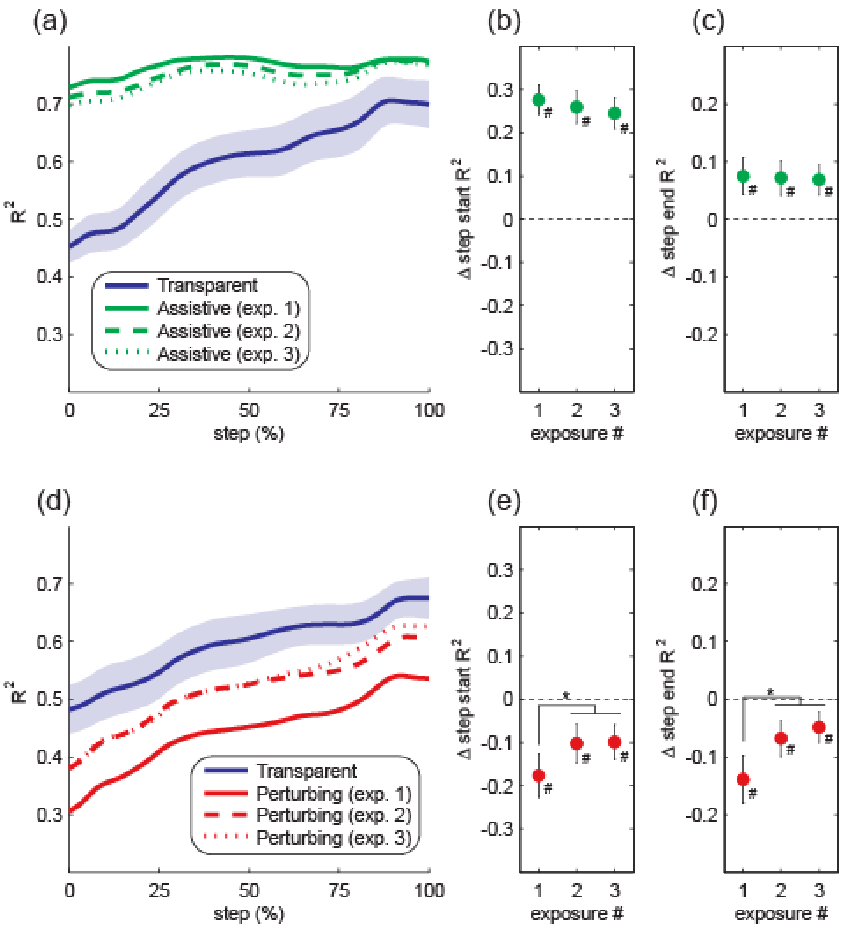


**Figure C3.** Effects of repeated exposure to the force-field. The top row (a-c) illustrates the direct effects of force field assistance, while the bottom row (d-f) illustrates the direct effects of force-field perturbations. Asterisks (*) indicate significant differences between the indicated exposures. Pound signs (#) indicate a significant difference from the initial Transparent trial.

*After-effects with repeated force-field exposure.* $R^2$ magnitudes throughout the step were similar during each of the washout periods following force-field assistance (Fig. C4a). No significant differences were present between these washout periods in terms of step start $R^2$ (p=0.86; Fig. C4b) or step end $R^2$ (p=0.85; Fig. C4c). During the washout periods following force-field perturbations, $R^2$ magnitudes throughout the step increased for later exposures (Fig. C4d). Both step start $R^2$ (p=0.048; Fig. C4e) and step end $R^2$ (p=0.002; Fig. C4f) increased significantly following later perturbation exposures.

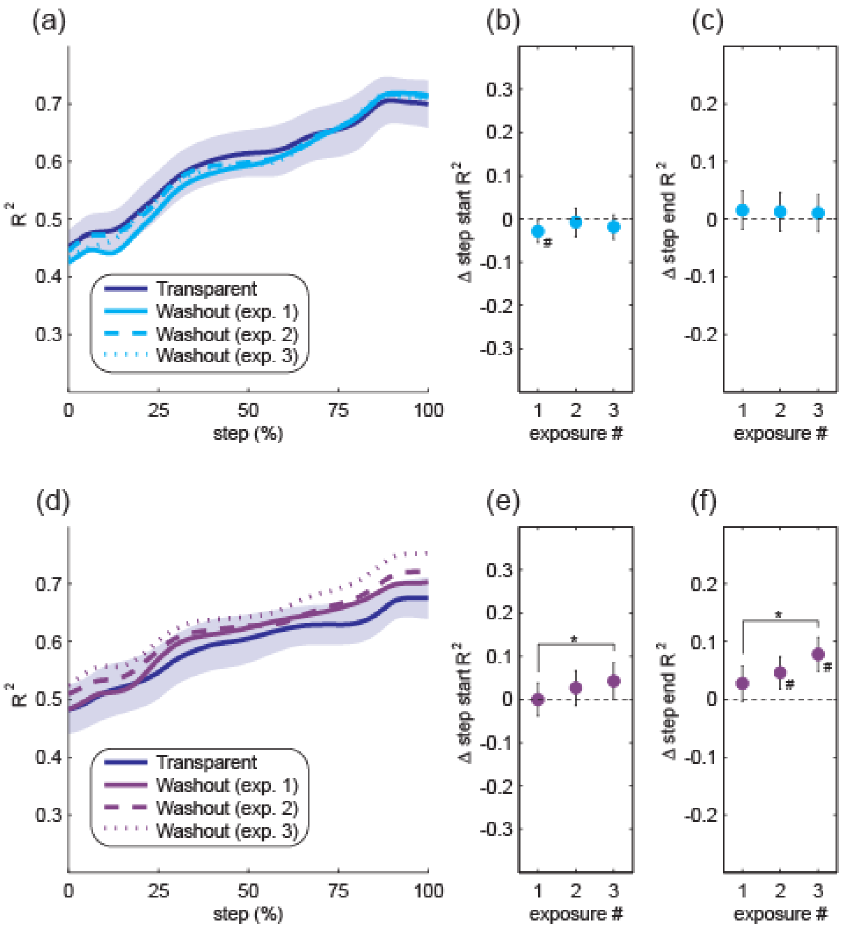


**Figure C4.** Gait behavior across three washout periods. The top row (a-c) illustrates gait metrics during the three washout periods following force-field assistance, while the bottom row (d-f) illustrates these metrics during washout periods following perturbations. The structure of the figure is the same as that described for Figure C3.

### Supplementary Information References

1. Nyberg, E.T., Broadway, J., Finetto, C., Dean, J.C. A novel elastic force-field to influence mediolateral foot placement during walking. *IEEE Trans. Neur. Syst. Rehabil. Eng.* **25**, 1481-1488 (2017).
2. Stimpson, K.H., Heitkamp, L.N., Horne, J.S., Dean, J.C. Effects of walking speed on the step-by-step control of step width. *J. Biomech.* **68**, 78-83 (2018).
3. Heitkamp, L.N., Stimpson, K.H., Dean, J.C. Application of a novel force-field to manipulate the relationship between pelvis motion and step width in human walking. *IEEE. Trans. Neur. Syst. Rehabil. Eng.* **27**, 2051-2058 (2019).
